# Deciphering Pre-existing and Induced 3D Genome Architecture Changes involved in Constricted Melanoma Migration

**DOI:** 10.1101/2024.08.21.609017

**Authors:** Christopher Playter, Rosela Golloshi, Joshua H. Garretson, Alvaro Rodriguez Gonzalez, Taiwo Habeeb Olajide, Ahmed Saad, Samuel John Benson, Rachel Patton McCord

## Abstract

Metastatic cancer cells traverse constricted spaces that exert forces on their nucleus and the genomic contents within. Cancerous tumors are highly heterogeneous and not all cells within them can achieve such a feat. Here, we investigated what initial genome architecture characteristics favor the constricted migratory ability of cancer cells and which arise only after passage through multiple constrictions. We identified a cell surface protein (ITGB4) whose expression correlates with increased initial constricted migration ability in human melanoma A375 cells. Sorting out this subpopulation allowed us to identify cellular and nuclear features that pre-exist and favor migration, as well as alterations that only appear after cells have passed through constrictions. We identified specific genomic regions that experienced altered genome spatial compartment profiles only after constricted migration. Our study reveals 3D genome structure contributions to both selection and induction mechanisms of cell fate change during cancer metastasis.

## Introduction

Tumors are highly heterogeneous and only a subset of cells within the tumor possess characteristics that make them more likely to successfully metastasize to secondary sites.^1,2^ Metastasis involves a multi-step cascade that includes invasion through dense extracellular matrix (ECM), intravasation into the bloodstream, and extravasation to a secondary site.^3^ During these steps, migrating cancer cells often must squeeze through spaces much smaller than the diameter of their nucleus, a process we refer to in this work as “constricted migration”. Cells with an initial increased propensity for constricted migration may therefore selectively end up at the metastatic site. However, other alterations in cell phenotype may occur during the process of metastatic migration, as a result of exposure to different microenvironments and the physical forces on the cells as they squeeze through tight constrictions. In some cases, metastatic tumors have been found to contain both “leader” and “follower”-type cells, in which the pre-existing cellular properties of the leaders are selected for, but further changes can occur in follower cells as they are exposed to new microenvironments^4^. These features of initial heterogeneity and environment-induced changes can be observed in cell culture models of cancer as well. As examples of heterogeneity, numerous different cancer cell lines in culture have been shown to contain stable subpopulations of stem-like cells^5,6^ or other subtypes that are marked by the expression of certain proteins.^7,8^ The A375 human melanoma cell line, which is the focus system in this study, has previously been found to exhibit heterogeneity and the potential for multiple subpopulations with different metastatic capacities in mice^9^ and in vitro migration experiments^10^. In addition to cellular differences created by this initial cellular heterogeneity, external stresses such as physical forces applied on cells or alterations to substrate stiffness can further alter cellular phenotypes and epigenetic marks.^11–13^

For better diagnostics and treatment, it is key to develop an understanding of which cellular features pre-exist and increase a cell’s initial propensity to migrate and to distinguish these factors from further changes that happen during metastatic migration and influence the cancer cell’s aggressiveness at a secondary site. Distinguishing these features is difficult, however, given the complex combination of selection from a heterogeneous population as well as induced alterations that occur in metastasis as discussed above. Emerging evidence has identified nuclear architecture and 3D genome structure as a potential key factor in cancer progression and metastasis. Genome wide chromosome conformation capture (Hi-C) experiments across multiple cancer types have identified changes in spatial genome organization patterns that correlate with cancer progression, epithelial to mesenchymal transition, and metastasis.^14–18^ In particular, alterations in cancer phenotype are often associated with changes in the spatial compartmentalization of chromosome regions into active (A) and inactive (B) compartments.^19^ However, these previous studies have not been able to determine which of these “progression associated 3D genome changes” could have pre-existed in a subpopulation of cells and been selected for and which changes could be induced during the process of metastasis.

Many questions also remain about how migration-favoring initial states or induced alterations are encoded. There is evidence that both initial heterogeneity and changes induced by stresses could be encoded by patterns of gene expression and epigenetic chromatin modification changes. Subtypes of breast cancers and B cell neoplasms are characterized by differences in 3D genome structure^18,20^. Computational models suggest that 3D chromosome folding can help encode stable memories of epigenetic states^21^, potentially maintaining subpopulation identity across cell generations. Meanwhile, in terms of induced changes, passage through multiple constrictions in a microfluidic device was shown to increase the histone modification H3K27me3 and decrease H3K9ac in HT1080 cells.^22^ Here, we investigate both what chromosome structures are associated with initial migration ability, and which are specifically changed only after passage through constrictions.

Previously, we have found that human melanoma cells (A375) that have passed through ten rounds of constricted migration exhibit stable differences in migratory phenotype, gene expression, nuclear architecture, and 3D genome structure^10^. Our previous results could not fully distinguish whether these changes were the result of selection on an initially heterogeneous population or the result of constriction-induced changes. We found that new populations of cells grown from individual clones of the parental A375 cell population underwent similar changes after sequential constricted migration. However, it is known that cancer cells can regenerate heterogeneity as they divide^23^, so the changes with constriction migration in the clonal populations could still be in part due to selection. Here, we identify a cell surface protein, Integrin-β4 (ITGB4), as a marker whose RNA and protein expression level closely associates with A375 constricted migration proficiency. ITGB4 is the beta subunit of a heterodimeric receptor for extracellular matrix components and select cell adhesion molecules with the preferred binding partner Integrin Alpha-6^24^. This protein has often been discussed in previous literature for its potential role in metastatic propensity, though depending on the cancer type and context, high ITGB4 levels may be a migration-favoring^25–28^ or inhibiting factor^29^. Overall, an increase in ITGB4 expression tends to lead to an increase in migration potential for most cancers^29^. ITGB4 is currently used as a prognosis marker for both pancreatic^25^ and lung cancers^30^. ITGB4’s role in melanoma is less well understood, but some results have indicated that an increase in its expression has led to higher migration abilities^31^. Indeed, we find that cells within the Parental A375 population with higher ITGB4 expression (ITGB4(+)) initially migrate at a higher efficiency through 5µm pores, a pore size that substantially constricts the cell nucleus. By isolating ITGB4(+) cells from the parental A375 population, we can identify pre-existing cellular characteristics and 3D genome structures that favor migration. Additionally, we find key differences between ITGB4(+) cells and cells that have passed through 10 rounds of constriction (Bottom10). By using this model, we can tease apart pre-existing 3D genome structures that enable constricted migration and 3D genome structures that arise as a result of constricted migration.

## Results

### ITGB4 expression is increased following A375 constricted migration and is associated with higher initial migration rates

Our previously published RNA-seq data^10^ for A375 cells that had passed through 10 rounds of 5µm constrictions revealed over 1000 differentially expressed (DE) genes following constricted migration (Bottom10) when compared to unmigrated cells (Parental) **(Figure 1A).** Of the most upregulated genes, Integrin-β4 (ITGB4), in particular stood out as a potentially useful cell surface marker that would enable sorting and tracking of cells and as a protein which had previously documented involvement in increasing cancer metastasis. We found that this upregulation occurs in constricted cells only, not when A375 cells migrate through 12µm pores that exert much less deformation on the nucleus **(Figure 1B).** Thus, increased ITGB4 gene expression cannot be an effect of one of the other factors common to migration such as fibronectin or serum gradient exposure. We further found that surface protein levels of ITGB4, as measured by flow cytometry (see Methods), increased after 10 rounds of constricted migration not only in the Parental population, but also in A375 populations grown from four isolated individual cells (Clonal populations 1-4) **(Figure 1C).** This shows that rather than an ITGB4 expressing cells only being a static subset of the parental A375 cell line, high levels of this protein can result from constricted migration even in different clonal lines. Since ITGB4 is expressed on the cell surface, we were able to use flow cytometry to sort out cells that highly expressed ITGB4 from the Parental population, resulting in a subgroup we term ITGB4(+), as well as cells that had almost no ITGB4 surface expression, which we call ITGB4(-). Using fluorescence activated cell sorting (FACS) gating based on an unstained control (see Methods), we isolated ∼20% of cells as ITGB4 expressing (ITGB4(+)) in the Parental A375 population while ∼35% expressed very little or none of this protein (ITGB4(-)) **(Figure1D).** The ITGB4(+) cells sorted out from the Parental population maintained high ITGB4 expression over rounds of cell division, even across 7 weeks of growth **(Supp Fig. 1A).**

**Figure 1.**
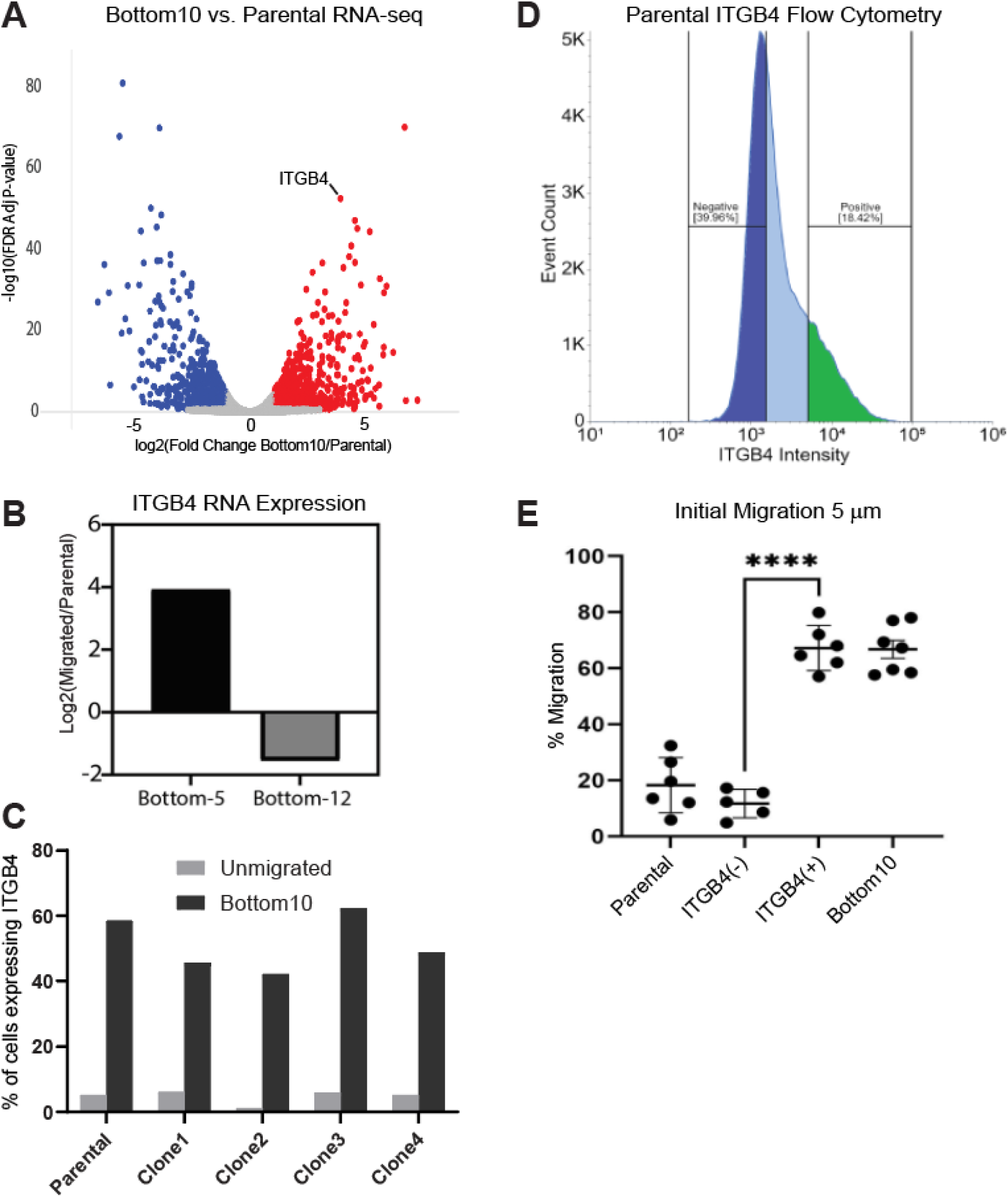
ITGB4 expression is increased following A375 constricted migration and is associated with higher initial migration rates. **A)** RNA-seq analysis of all genes comparing Bottom10 vs. Parental cells. Significantly upregulated (red) or downregulated (blue) and unchanged (grey) genes are shown. ITGB4 labeled. **B)** ITGB4 log2(Migrated/Parental) RNA levels after 10 rounds migration through 5µm (constricting) and 12µm (non-constricting) pores. **C)** Flow cytometry measured percentage of cells that express surface ITGB4 protein for Parental and all 4 Clonal populations before migration and after 10 rounds of sequential migration. **D)** ITGB4 intensity plot (based on green fluorescent antibody labeling) used to FACS sort ITGB4(-) cells (*dark blue)* and ITGB4(+) cells (*green).* Cells in the light blue section were not collected. **E)** The initial migration rates (% of cells that migrated through pores) through 5µm Transwell pores for Parental, ITGB4(-), ITGB4(+) cells along with the final migration rate for Bottom10 cells. (****P=<0.0001, two-tailed t-test)

We next challenged these FACS-sorted ITGB4(+) and ITGB4(-) cells to migrate through 5μm pore Transwell filters to test their initial migratory ability. We found that the ITGB4(+) cells migrated at a significantly higher rate than ITGB4(-) cells **(Figure 1E).** In fact, the ITGB4(+) cells migrated about as well as Parental cells after 10 rounds of constriction (Bottom10). This raises the question: is the ITGB4(+) sub-population the group that is being selected by rounds of constriction and thus isolated as Bottom10 cells at the end of sequential constrictions? Thus, we further characterized ITGB4(+) phenotypes in comparison to ITGB4(-) and Bottom10 cells.

### ITGB4(+) cells exhibit phenotypic differences that represent an intermediate between ITGB4(-) and sequentially constricted Bottom10 cells

Cells that express ITGB4 exhibit morphological differences compared to ITGB4(-) cells. ITGB4(-) cells had a more epithelial morphology, whereas ITGB4(+) cells exhibited fewer cell-cell adhesions and were more likely to extend reaching protrusions similar to Bottom10 cells **(Figure 2A).** However, the growth of these cell populations on a 2D surface also provided a first indication that ITGB4(+) cells were not quite the same as Bottom10 cells. Quantifying the percentage of cells with extended protrusions showed that ITGB4(+) cells had a significantly higher percentage of cells with protrusions compared to ITGB4(-) cells, but not as many as Bottom10 cells **(Figure 2B).** The cell body aspect ratio across the imaged cells is similar between ITGB4(+) and ITGB4(-) and both were significantly less elongated than Bottom10 cells **(Figure 2C)**. Nonetheless, ITGB4(+) cells showed increased wound healing in a 2D scratch assay compared to the ITGB4(-) cells, which showed very little wound closure even after 24 hours **(Figure 2D, Movies 1 and 2).** After seeing the differences between these cell groups in a 2D environment, we next asked if they behaved differently when embedded in a 3D collagen matrix. Cells were embedded in a 3D collagen matrix (see Methods) and allowed to acclimate for 96 hours. Again, there were visually apparent differences between ITGB4(-) and ITGB4(+) cells **(Figure 2E).** ITGB4(-) cells remained spherical and failed to migrate through the matrix, instead rotating in place. ITGB4(+) cells, however, grew extremely long protrusions through the collagen matrix. These protrusions were similar to those exhibited by Bottom10 cells placed in a collagen matrix, but while Bottom10 cells also deformed their nuclei to translocate through the matrix, ITGB4(+) cell nuclei remained largely spherical and did not translocate through the dense matrix **(Figure 2E, Movies 3-4 (ITGB4-), 5-6 (ITGB4+), 7 (Bottom10))**.

**Figure 2.**
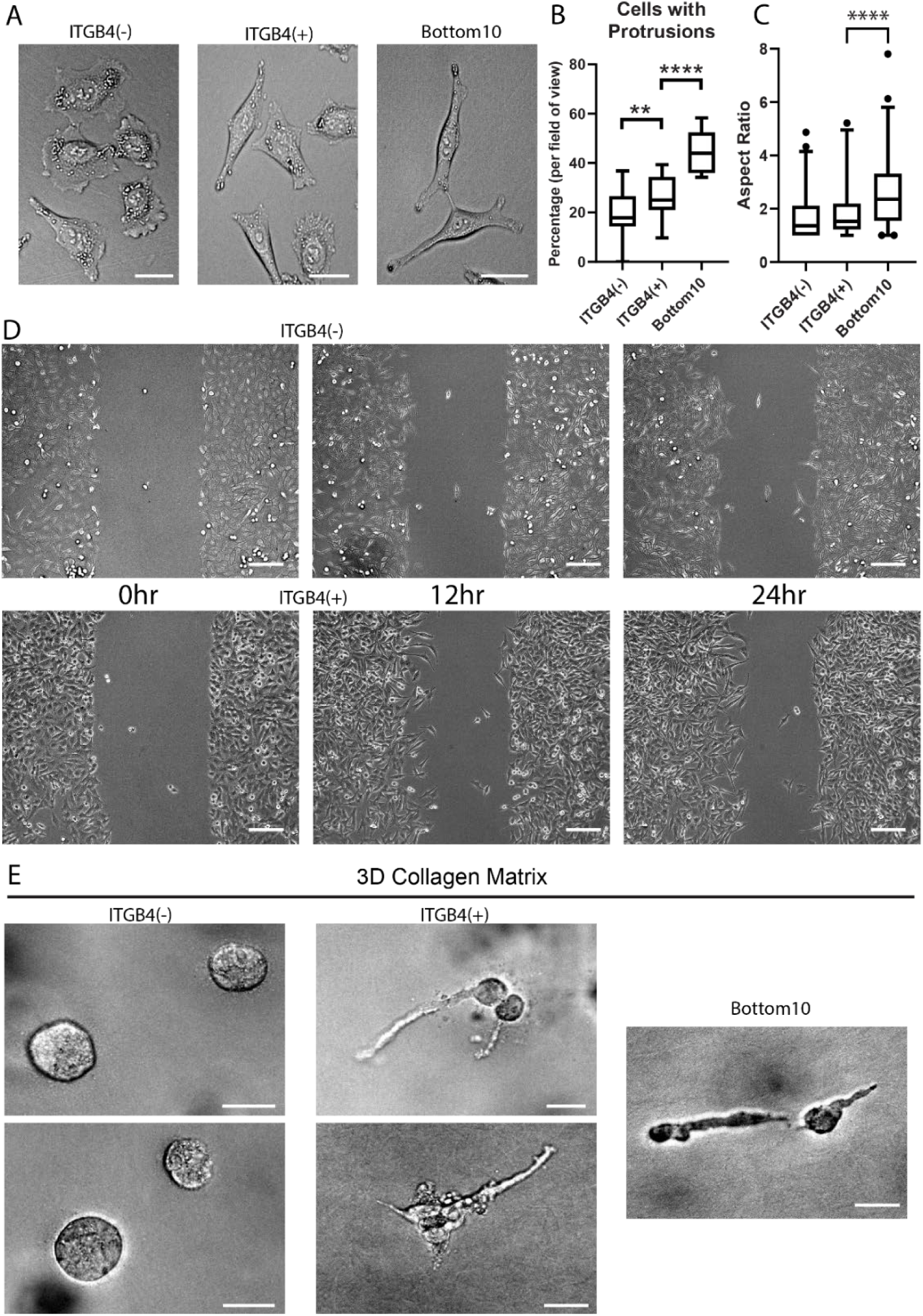
ITGB4(+) cells exhibit phenotypic differences that represent an intermediate between ITGB4(-) and sequentially constricted Bottom10 cells. **A)** 2D phase contrast images of ITGB4(-), ITGB4(+), and Bottom10 A375 cells. Scale = 70 µm. **B)** Quantification of the percentage of cells that have protrusions for ITGB4(-), ITGB4(+), and Bottom10 cells per field of view. Images taken at 20x on EVOS. Total cells counted: 695-ITGB4(+), 559-ITGB4(-), 468- Bottom10. (**P=0.0048, ****P<0.0001, two-tailed t-test) **C)** Aspect ratio quantification for ITGB4(-), ITGB4(+), and Bottom10 A375 cells. (****P<0.0001, two-tailed t-test) N=∼100 cells/condition **D)** Scratch Assay images (20x, phase contrast) at 0, 12, and 24hr post scratch for both ITGB4(-) and ITGB4(+) cells. See Movies 1 and 2. Scale = 300µm. **E)** 40x Phase contrast images of ITGB4(-), ITGB4(+), and Bottom10 cells embedded in 3D collagen matrix 96hr post seeding. See Movies 3-7. Scale = 70µm.

### ITGB4 expression increase across successive rounds of migration shows both selection and induction features

To further investigate to what extent sequential constriction selects for high ITGB4 expressing cells due to their initially high migration efficiency, we next monitored in detail the levels of ITGB4 cell surface expression across the rounds of constricted migration, starting with the parental cell population. If cells expressing ITGB4 are enriched by each round of migration, we would expect an increase in the percentage of ITGB4 expressing cells found immediately after each round of migration compared to just before migration. Using flow cytometry, we determined the percentage of cells that highly expressed ITGB4 before and after each round of 10 sequential rounds of migration through 5μm Transwell pores **(Figure 3A).** This experiment allowed us to monitor ITGB4 expression in 3 different settings: constricted migration (5μm), non-constricted migration (12μm), and no migration at all (No Well).

**Figure 3.**
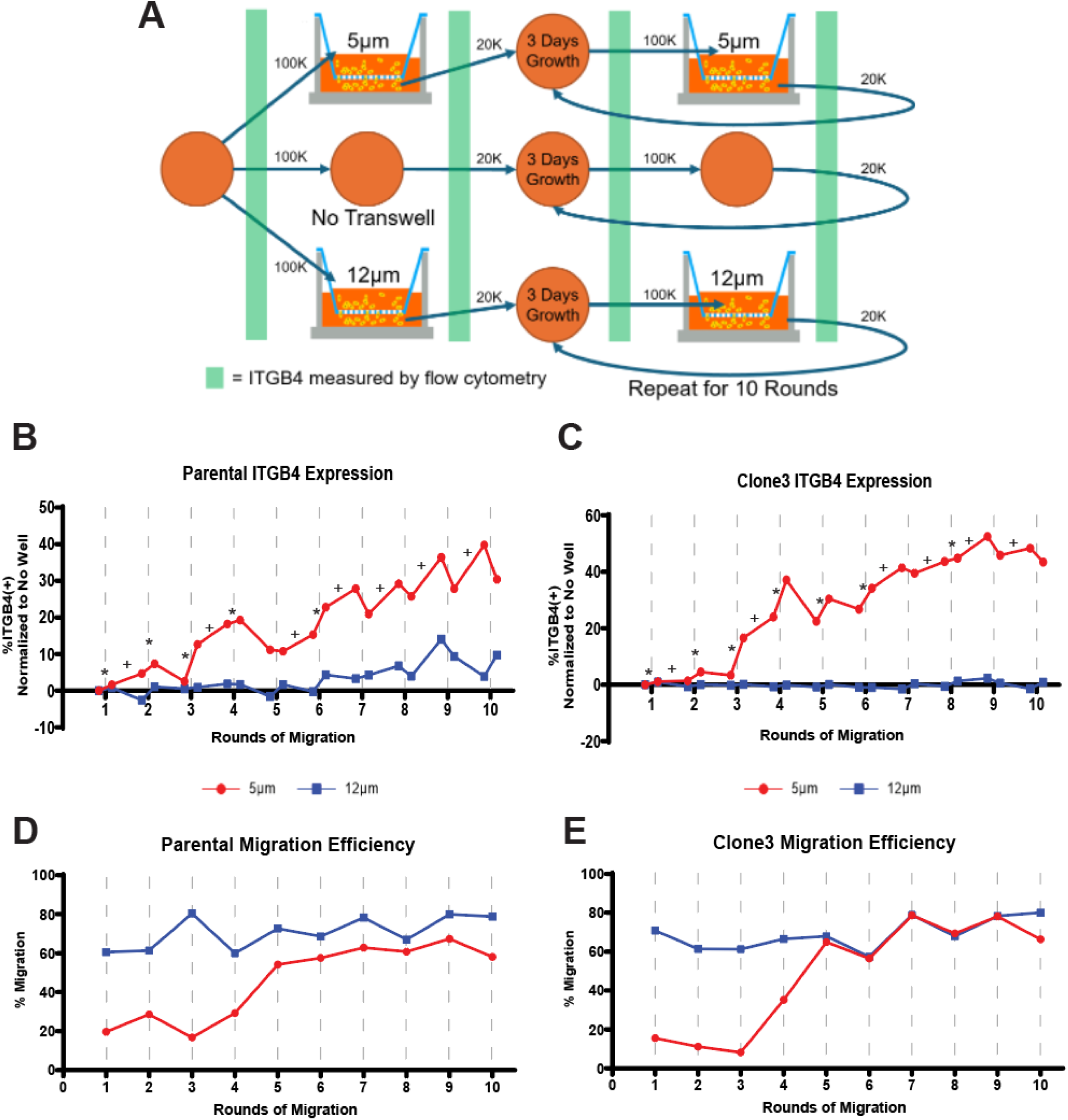
ITGB4 expression increase across successive rounds of migration shows both selection and induction features. **A)** Experimental approach to monitor ITGB4 surface expression across rounds of constricted migration. Orange circles represent culture dishes. For all conditions, 100K cells were always used to start the Transwell experiment, and 20K cells taken for the subsequent 3 days growth. **B)** ITGB4 surface protein expression (flow cytometry) over sequential rounds of migration for 5µm, *red,* and 12µm, *blue,* Transwell pores normalized to the No Well condition. Vertical dashed lines represent a single Transwell migration event. Stars indicate selection-like increase after Transwell passage, while crosses indicate further induction during growth. **C)** Clone3 ITGB4 surface protein expression (flow cytometry) over sequential rounds of migration. All colors and symbols as in (B). **D)** Parental migration rates (% of cells that migrated through pores) for 5µm, *red*, and 12µm, *blue*, pores throughout the 10 rounds of sequential migration matching the flow cytometry data conditions in (B). **E)** Clone3 migration rates matching the 10 rounds of sequential migration matching the conditions in (C).

Previous work has shown that cell surface integrin expression can be influenced by cell growth conditions such as confluence and starvation.^32,33^ We found that in A375 cells, ITGB4 protein expression on the cellular surface was increased additively by both cell confluency and media depletion **(Supp. Fig. 1B-D)**. Therefore, we carefully normalized our ITGB4 expression results across sequential migration by the “No Well” control, which mimicked the same confluence and media conditions at all timepoints, but without any migration. During the sequential migration, both the Parental population and the cells derived from a single A375 clone (Clone3) showed an increase in ITGB4 surface protein expression level as they progressed through more rounds of migration **(Figure 3B and C).** Importantly, the increase was specific to constricted migration (not observed in the 12µm pore experiment), indicating that other conditions such as exposure to fibronectin did not cause the increase. In some rounds of migration (see stars in Figure 3B and C), we saw the expected signature of selection, where ITGB4(+) cells increased immediately after cells were passed through the Transwell. However, in other rounds of migration, this was not the case: instead, cells increased ITGB4 levels during growth after migration (see crosses, Figure 3B and C). This provides evidence of both selection from the previous population and some other mechanism of further ITGB4 induction after constricted migration.

Across rounds of sequential migration, the migration rate at each round correlates well with the ITGB4 expression both before the Transwell (0.823, 0.927, Pearson’s R) and the ITGB4 expression measured immediately after migration (0.770, 0.874) for Parental and Clone3 cells respectively **(Figure 3D and E)**. Specifically in the round 3-5 range, both ITGB4 expression and migration ability exhibit large increases. Again, if passage through the Transwell filter simply selected for ITGB4 expressing cells, we would expect that the 20% of cells that migrate in the first round **(Figure 3D and E)** would all have high levels of ITGB4 expression. Instead, we see that it actually takes several rounds of migration to produce the substantial increase in both ITGB4 expression and migration efficiency. Together, the data suggest that ITGB4 can be an initial selection marker for migratory cells, but can also be further increased in the cell population as a result of migration.

### Knockdown of ITGB4 alters migration ability of previously unmigrated cells

The strong correlation between ITGB4 expression and constricted migration proficiency raised the question: is ITGB4 presence on the cell surface needed for effective constricted migration? To test this, we knocked down the expression of ITGB4 using RNAi in both ITGB4(+) and Bottom10 cells. By flow cytometry, we found that RNAi decreased cell surface ITGB4 expression by 60% in both cell populations **(Figure 4A and C)**. However, we found that this ITGB4 reduction affected constricted migration rates only in the previously unmigrated ITGB4(+) cell population. In these cells, reducing ITGB4 levels decreases constricted migration rates moderately, but significantly **(Figure 4B, Sup.** Fig. 2). In contrast, the Bottom10 cells, which also have high ITGB4 levels initially, but have already passed through 10 rounds of constriction, exhibited no migration difference following knockdown of ITGB4 expression **(Figure 4D, Sup.** Fig. 2). Thus, ITGB4 expression is more important to the migration ability of cells that have not previously passed through constrictions.

**Figure 4.**
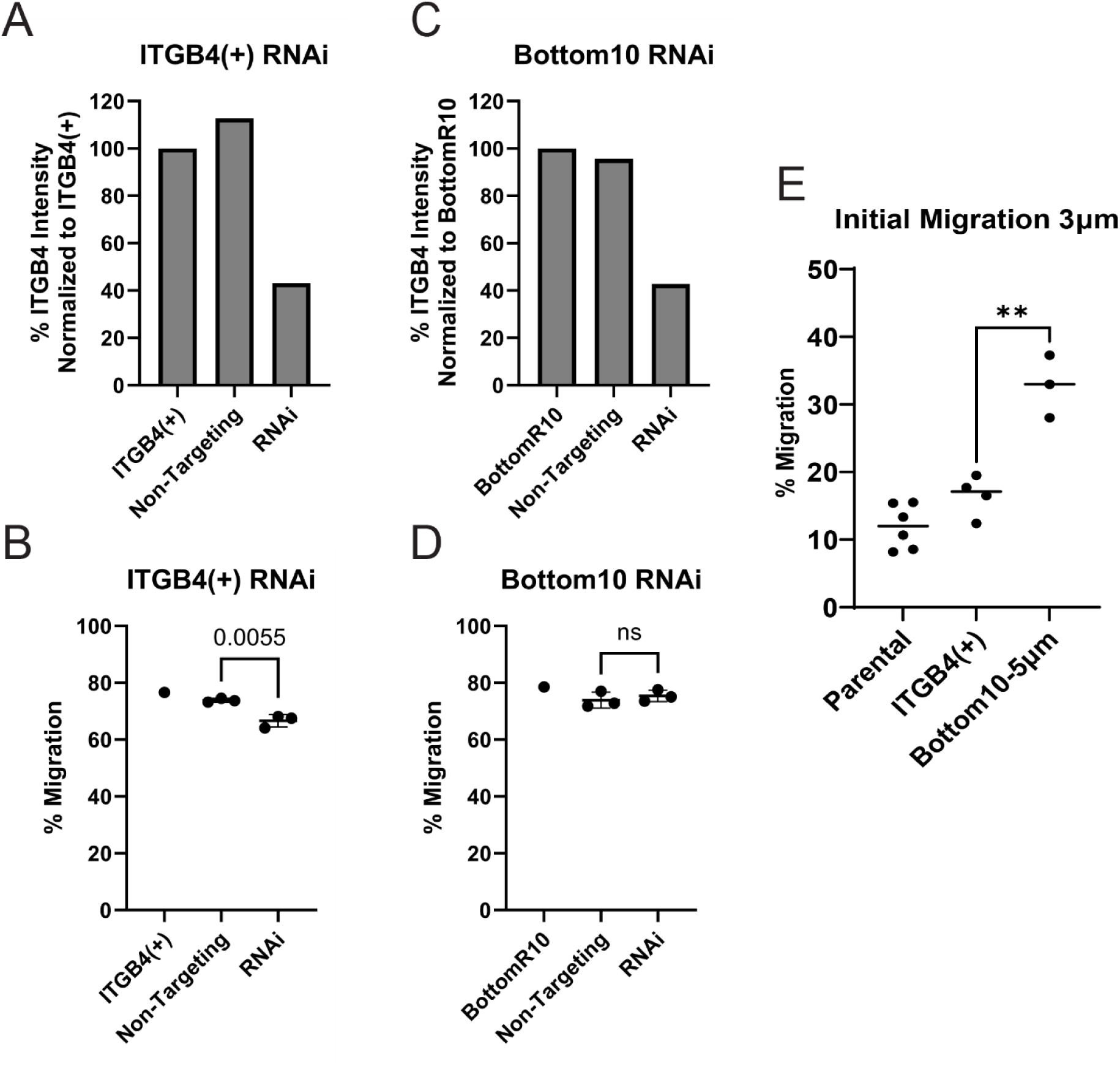
Knockdown of ITGB4 alters migration ability of previously unmigrated cells. **A)** Bar plot quantifying ITGB4 surface protein expression levels (flow cytometry) for untreated ITGB4(+) cells compared to those given non-targeting control RNAi, and ITGB4 RNAi for 96 hours. **B)** Migration rates (% of cells that migrated through 5μm pores) for conditions in Figure 4A. Technical replicates for non-targeting and ITGB4 RNAi shown (two tailed t-test). Other biological replicates are shown in Sup. Figure 2. **C)** ITGB4 surface protein expression in control and RNAi knockdown of Bottom10 cells, conditions as in (A). **D)** Migration rates for conditions in Figure 4C through 5µm Transwell pores. Technical replicates for non-targeting and ITGB4 RNAi shown. Other biological replicates are shown in Sup. Figure 2 (two tailed t-test). **E)** Initial migration rates for Parental, ITGB4(+), and Bottom10 cells when challenged with 3µm Transwell pores. (**P=0.0024, two tailed t-test) Biological replicates shown.

Combined with the phenotypic differences observed between ITGB4(+) and Bottom10 cells in Figure 2, this result points to additional phenotypic changes that happen as a result of sequential constricted migration that are not present in an initially highly migratory subpopulation. Further supporting this idea, we found that while ITGB4(+) cells migrated through 5μm pores at similar rates as Bottom10 cells, when we challenged both cell groups to migrate through even smaller 3μm pores, we found that Bottom10 cells migrated at significantly higher rates than ITGB4(+) cells **(Figure 4E)**. This suggests that having experienced the stress of 5μm pores aided the Bottom10 cells’ ability to pass through the tighter constrictions, which involve even greater nucleus deformation.

### 3D Genome structure differences exist between ITGB4(+) and ITGB4(-) cells at the compartment level

Based on the combination of migration ability and the morphological differences we saw between our ITGB4(+) and ITGB4(-) cells, we next asked if they also had altered 3D genome structures that associated with such differences. High throughput chromosome conformation capture (Hi-C) was carried out on both cells sorted for high and low ITGB4 expression in the Parental and one Clonal population. Many features of genome structure are preserved between these different cell groups, as seen by visual inspection of chromosome contact maps **(Figure 5A and B)**. Difference maps emphasizing changes in contact frequency reveal areas of contact change that indicate differences in the plaid A/B compartmentalization pattern. Such changes typically indicate regions that alter their spatial organization to associate more with heterochromatic (B) or euchromatic (A) regions in the nucleus. When looking at comparison heatmaps for both Parental **(Figure 5A)** and Clone3 **(Figure 5B)** on Chr17, we saw that the differences between ITGB4(-) and ITGB4(+) cells occurred in similar locations. This difference is easier to see when plotting the principal component analysis first eigenvector values for each 250kb region of Chr17. This compartment analysis shows genomic regions where both the Parental ITGB4(+) and Clone3 ITGB4(+) cells exhibit a strong shift towards the B compartment **(blue box, Figure 5C)**. To see how many genomic regions shift towards the A compartment or B compartment on a more global scale, we compiled every 250kb region that had a 20% shift in eigenvector value in either direction and plotted them on a phenogram^34^ **(Figure 5D).** This phenogram then showed us the position and direction of every compartment shift for both Parental and Clone3 cells when comparing ITGB4(-) to ITGB4(+) cells. While there are hotspots where shifts occur more frequently, nearly every chromosome had some region that shifted compartments. We found many areas where Parental and Clone3 shifts were concordant, and though not all changes were observed in both cell groups, no regions changed in opposite directions in Parental vs. Clone3 ITGB4(+) cells. This indicates that, overall, there is a consistent 3D genome compartment pattern that characterizes the initially migratory ITGB4(+) subpopulation.

**Figure 5.**
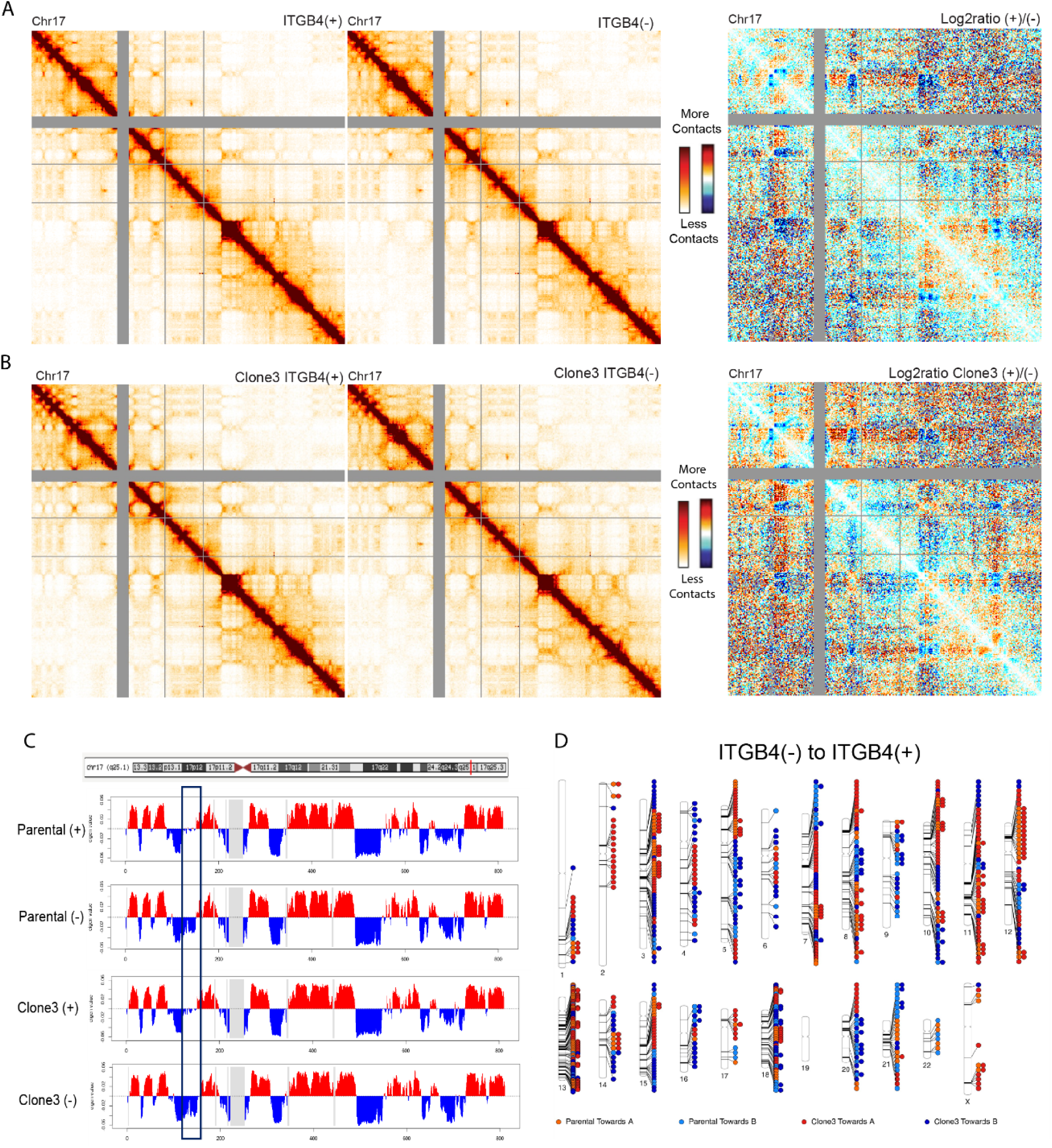
3D Genome structure differences exist between ITGB4(+) and ITGB4(-) cells at the compartment level. **A)** 250kb binned Hi-C interaction heatmaps of chr17 for ITGB4(+) and ITGB4(-) cells, along with the log2ratio of the conditions (blue = fewer contacts in ITGB4(+) cells compared to ITGB4(-) cells, red = increased contacts in ITGB4(+) cells). **B)** 250kb binned Hi-C interaction heatmaps of chr17 for Clone3 ITGB4(+) and Clone3 ITGB4(-) cells, along with the log2ratio of the conditions, as in (A). **C)** PC1 compartment profile for chr17 for ITGB4(+) and ITGB4(-) cells from both Parental and Clone3 populations. Dark blue box highlights a region where both Parental and Clone3 ITGB4(+) cells differ from ITGB4(-) cells. The red line in the chromosome ideogram marks the location of the ITGB4 gene on chr17 (always in a strong A compartment). **D)** Phenogram showing all compartment shifts (change in eigenvector1 value > 0.04 for Towards A or <-0.04 for Towards B) between ITGB4(-) and ITGB4(+) cells for both Parental and Clone3 populations. Each circle represents a single 250kb bin that is shifted, and its location is marked by a line to its respective chromosome. Regions with 2 circles side by side show shifts found in both populations.

### The 3D genome structure of ITGB4(+) cells is an intermediate state between Parental and Bottom10 cells

Morphologically and functionally, our results thus far have revealed the ITGB4(+) cells from the unmigrated population share some but not all characteristics of sequentially migrated Bottom10 cells. Our observation that ITGB4(+) cells exhibit 3D genome compartment differences from ITGB4(-) next led us to ask whether these are the same alterations we had previously observed in Bottom10 cells after sequential migration^10^. Therefore, we used our Hi-C data to identify regions of the genome where ITGB4(+) cells had the same compartment identity as Bottom10 cells **(Figure 6A green box)** or the same compartment identity as unmigrated Parental cells **(Figure 6A purple box).** Finding examples of similarities to both Parental and Bottom10 cells suggested that, while some features of the ITGB4(+) genome structure before migration could be selected for and thus observed in Bottom10 cells, there were other portions of the genome structure that changed only after constricted migration. Genome-wide, we found 669 genomic bins (250kb resolution) that were shifted between Parental cells and Bottom10 cells.

**Figure 6.**
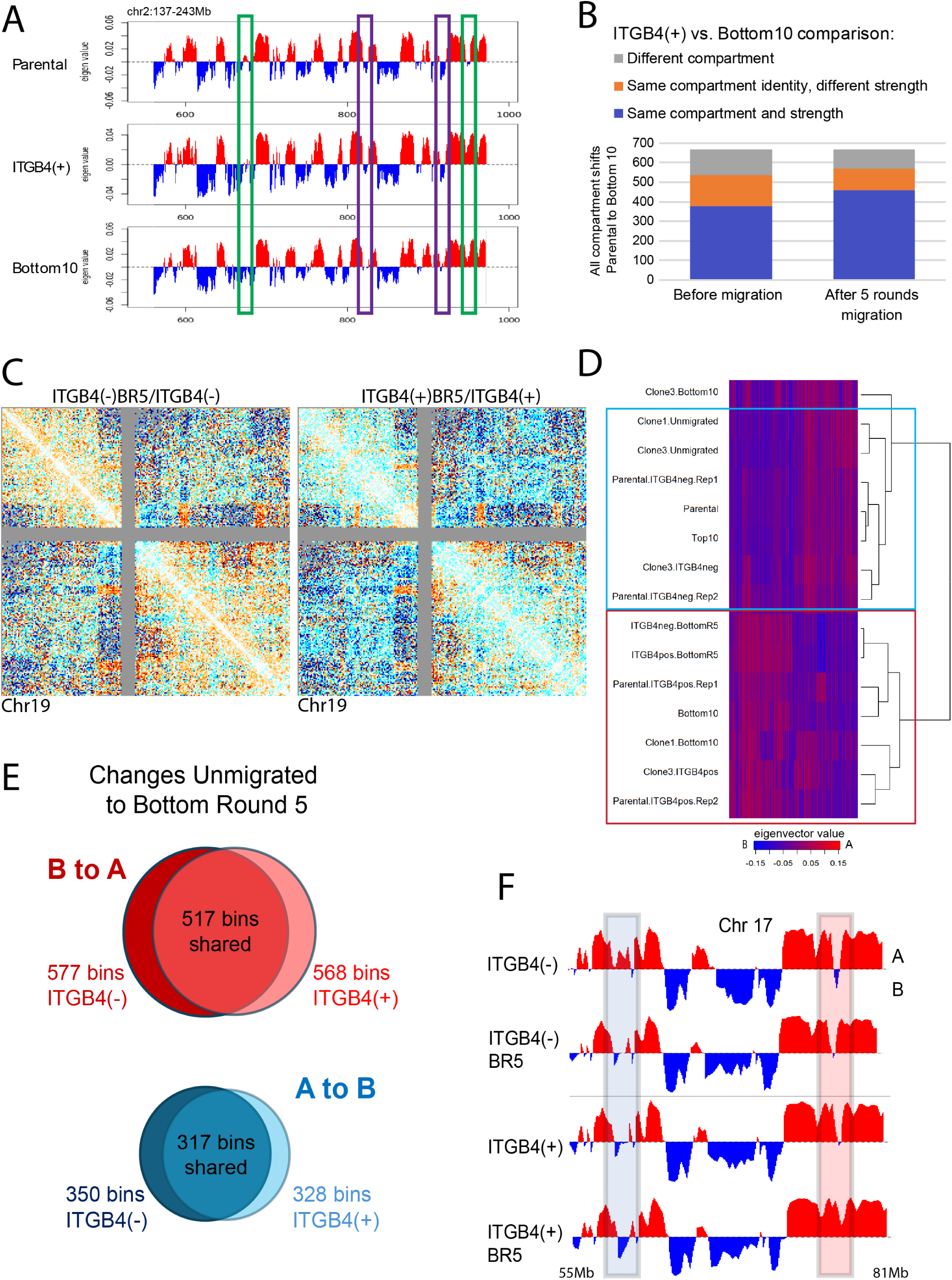
The 3D genome structure of ITGB4(+) cells is an intermediate state between Parental and Bottom10 cells. **A)** PC1 compartment profile for chr2:137-243Mb for Parental, ITGB4(+), and Bottom10 cells. Green boxes highlight regions where the compartment identity for ITGB4(+) cells is similar to Bottom10 cells, but different from Parental. Purple boxes show regions where ITGB4(+) cell compartment state is similar to Parental cells, but different from Bottom10 cells. **B)** Considering all 250kb bins that change compartment from Parental to Bottom10 cells (669 bins), colored bars indicate how many have matching compartment state and strength between ITGB4(+) and Bottom10 (blue), matching compartment state, but weaker in ITGB4(+) cells (orange), or ITGB4(+) cells match Parental rather than Bottom10 (grey). The left bar considers ITGB4(+) compartment state before any migration while the right bar considers ITGB4(+) cell compartment state after 5 rounds of migration. The proportion of ITGB4(+) compartments that match the strength and identity of Bottom10 before migration is high (blue = 379, left bar), but becomes even higher after these cells pass through 5 rounds of constriction (blue = 461, right bar). **C)** 250kb binned Hi-C heatmap showing the chr19 log2ratio of both 5 rounds migrated (BR5) ITGB4(-) and ITGB4(+) cells vs. unmigrated cells of those groups. **D)** Clustering analysis using the 200 most variable 250kb compartment bins. Light blue box: non-migratory cell conditions. Red box: migratory cell conditions. **E)** Analysis of 250kb regions that change compartment (- to + PC1 value for B to A, + to – PC1 value for A to B) for both ITGB4(-) and ITGB4(+) cells that migrated through 5 rounds of 5 µm pores. Outside values represent the total number of bins changed following migration; inner value is the number of those bins that changed in the same manner in 5 rounds of migration for both ITGB4(+) and (-). **F)** PC1 compartment profiles for chr17:55-81Mb for ITGB4(-), ITGB4(-)BR5, ITGB4(+), and ITGB4(+)BR5 conditions ordered from least to most migratory (top to bottom). Blue box= progressive shift in compartment identity towards a stronger B compartment as migration ability increases. Red box = progressive shift towards a stronger A compartment as migration ability increases.

Of those 669, 536 bins (80%) were in the same compartment in ITGB4(+) cells as in Bottom10 cells **(Figure 6B, left)**. There is a wide range of values that count as being in the “same compartment”, even though the compartment strength may be dissimilar, so we added another level of specificity to find that 379 bins (57%) in the ITGB4(+) cells were in the same compartment and within 10% of the compartment strength of Bottom10 cells.

### ITGB4(+) cells are sensitive to constricted migration and resemble Bottom10 cells more closely following multiple rounds of migration

From our comparisons above, we hypothesized that the remaining bins that were altered in Bottom10 vs. Parental, but not in ITGB4(+) cells, may arise from the process of constriction. To test that hypothesis, we took both ITGB4(-) and ITGB4(+) cells and allowed them to migrate through 5 rounds of 5μm constrictions **(Sup.** Fig. 1E). From the Hi-C comparison heatmaps **(Figure 6C)** we saw that the genome structure of both ITGB4(-) and ITGB4(+) cells were sensitive to migration and exhibited similar compartment changes afterwards. We then quantified compartment shifts as above and found that after ITGB4(+) cells had passed through 5 rounds of constriction, 568 bins (85%) were now in the same compartment as Bottom10 cells, and 461 bins (69%) were in the same compartment and within 10% of compartment strength **(Figure 6B, right).** This suggests that the act of passing through constrictions plays a pivotal role in altering the 3D genome even of already highly migratory cells and that the final 3D genome state of cells after sequential migration involves both selecting for properties already inherent to certain subpopulations of cells and further inducing alterations during the rounds of migration.

We clustered all cellular conditions according to compartment signatures **(Figure 6D).** In general, migratory cell conditions (red box) clustered together and separately from non-migratory conditions (light blue box). Notably, the most similar compartment profiles were found between the ITGB4(+) and ITGB4(-) cells that had migrated through 5 rounds of constriction.

This suggests that even though ITGB4(-) and (+) cells start out with distinct compartment profiles, constricted migration shifts their compartment profiles to become very similar. Indeed, when we identified bins that switch compartments after 5 rounds of migration, we found that about 90% of the constriction-associated switched regions are shared between ITGB4(+) and ITGB4(-) cells **(Figure 6E)**. We found that if we arrange conditions from least to most migratory (from poorly migratory ITGB4(-) cells to ITGB4(+) cells that had been migrated 5 times), we can see genomic regions which progressively shift compartments **(Figure 6F).** Shown in the blue box is an example where a strong A compartment shifts to a strong B compartment as the cells become more migratory. Just downstream, the red box highlights a shift in the opposite direction, where a strong B compartment shifts to becoming a strong A compartment. In each case, we observed that ITGB4(+) cells often already looked more similar to migrated ITGB4(-) cells, but then passage through constrictions of this already migratory population resulted in even further compartment shifts in the same direction as the migrated vs. unmigrated ITGB4(-). This trend was observed over many regions genome-wide. We identified regions that progressively shifted their compartment from ITGB4(-) to ITGB4(+)-BottomRound5 cells by calculating the slope of the eigenvector across these conditions and identifying regions where the slope (positive or negative) was greater than 1.5 standard deviations from the mean. This analysis revealed 668 regions that progressively shift towards A and 640 regions that progressively shift towards B (the top 30 most dramatic slopes are shown in **Figure 7A).**

**Figure 7.**
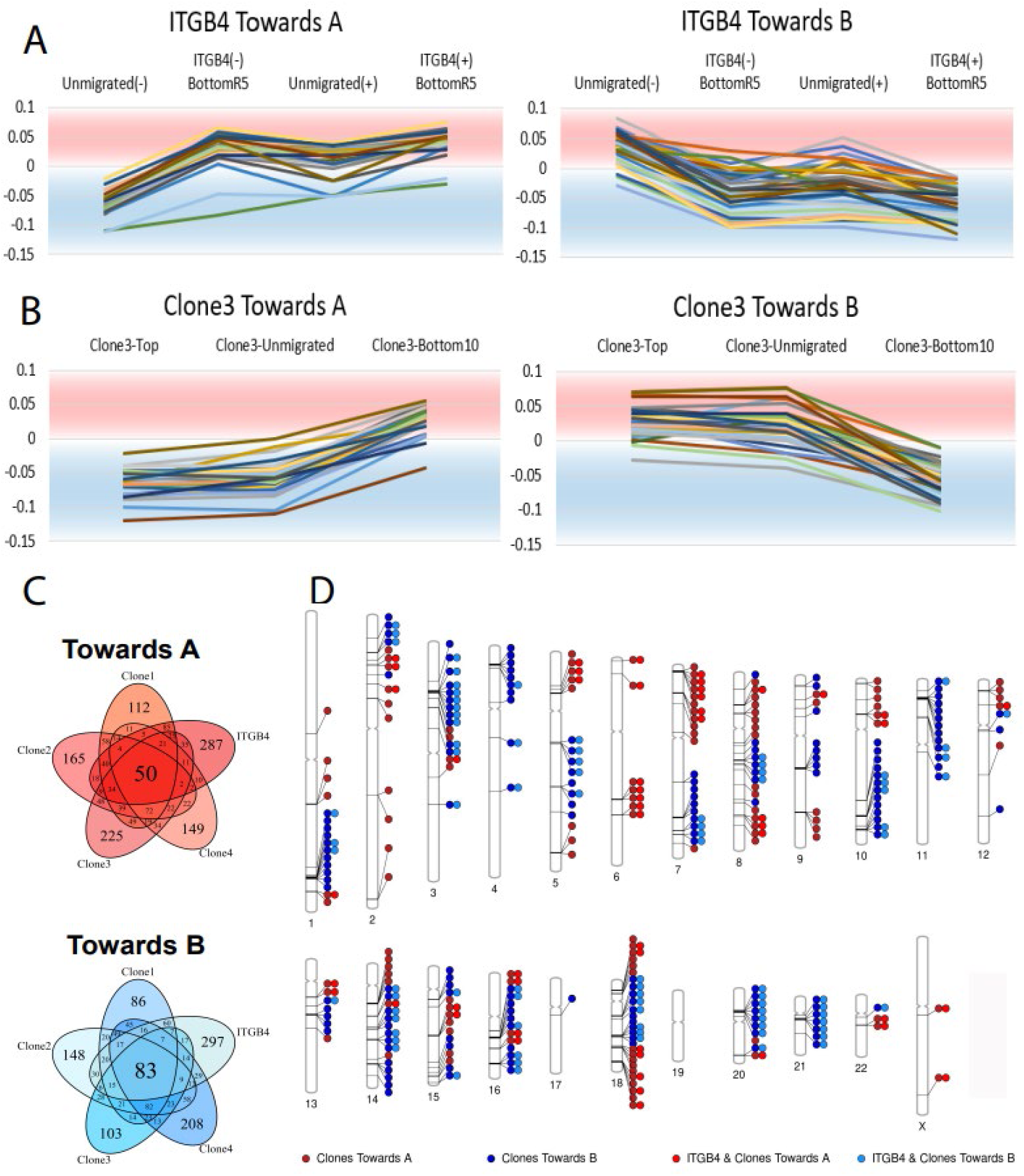
- Specific regions of the genome are prone to compartment shifts/switches following constricted migration. **A)** The top 30 250kb binned regions with positive slopes (Towards A) and negative slopes (Towards B) as cells increase migration ability. Each bin’s compartment value (PC1) is plotted for ITGB4(-), ITGB4(-)BR5, ITGB4(+), and finally ITGB4(+)BR5 and tracked from left to right. **B)** The top 30 250kb binned regions with positive slopes (Towards A) and negative slopes (Towards B) as cells increase migration ability for Clone3 cells. Each bin’s compartment value is plotted for Top10, Unmigrated, and Bottom10 cells and tracked from left to right. Other clonal populations can be found in Sup Fig 3. **C)** 5-way Venn diagram showing the number of 250 kb bins that shift Towards A or Towards B across either ITGB4 conditions or Clonal migration. All bins that have slopes +/- 1.5STD from the mean slope are included. The number on the outer most part of ovals represents the number of bins that are specific to each condition. The number in the center represents the number of bins that are shared across all 5 conditions. **D)** Phenogram showing locations of all shifted 250kb bins shared between the 4 Clonal populations (dark red and dark blue) and bins that are shared between the Clonal populations and the ITGB4 conditions (light red and light blue). Each circle represents a single 250kb bin that has shifted. Single circles show bins only changed across the 4 clonal populations, whereas double circles show regions where the ITGB4 conditions also shared that shift.

### Specific regions of the genome are prone to compartment shifts/switches following constricted migration

Our next goal was to determine the genomic regions that are the most susceptible to compartment shifts or switches following constricted migration. Previously, we had completed Hi-C on four different clonal populations at 3 different conditions: Top10 (unable to migrate though given 10 chances), Unmigrated, and Bottom10. We ordered these conditions from least to most migratory and using their eigenvector values for each 250kb region were able to calculate a slope for each region. In the same way that we did for the ITGB4 spectrum, we considered regions with a positive or negative slope greater than 1.5 standard deviations from the mean as shifted regions following constricted migration (top 30 most positive and negative sloped bins shown in **(Figure 7B and Sup.** Fig 3). Once we found which bins were the most susceptible to shifts for each clone and ITGB4 populations, we identified bins that overlap across all conditions. Across all clonal and ITGB4 conditions, 50 total 250kb bins shifted towards the A compartment and 83 total bins shifted towards the B compartment **(Figure 7C).** If we only consider the clonal populations those numbers increase to 122 bins towards A and 165 bins towards B. The shifts shared by clonal populations but not ITGB4 likely reflect the different initial starting state of ITGB4-sorted cells. We next investigated the genomic locations of these compartment shifts by plotting all the shifting regions on a phenogram **(Figure 7D).** We found that constriction-sensitive regions are located throughout the genome with all but chromosome 19 having at least one shifted region. Most chromosomes have regions that shift both towards the A and the B compartments. We observed hotspot areas, where specific chromosomes had larger regions that were sensitive, suggesting that these areas could be at play in enabling better migration, i.e., chromosome 3, 11, and 18. In particular, comparing chr17, chr18, and chr19, we note that chr18 experiences far more changes. Chromosomes 17 and 19 are gene rich and internally localized while chromosome 18 is gene poor and peripherally located. This suggests that chromosome gene density and localization may influence susceptibility to constriction-induced compartment switches. Overall, these data suggest that constricted migration can have an effect on 3D genome structure and that there are regions of the genome that are more susceptible to compartment shifts and switches than others.

### Gene expression changes partially correlate with compartment shifts following constricted migration

After identifying compartment shifts and switches in our Hi-C analysis, we next asked whether genes in those regions had altered gene expression. We performed RNA-seq on cell populations from a spectrum of ITGB4 and migration conditions and clustered the conditions based on their RNA expression profiles **(Figure 8A).** Similar to the compartment profiles, migratory cells clustered together and separately from non-migratory cells based on RNA profiles. The analysis found 6 gene clusters that best accounted for the variance with the most notable being the lime cluster, upregulated in migratory cells, and the light blue cluster, upregulated in the non-migratory cluster. Enrichr^35^ analysis of the genes within the lime cluster showed us that multiple pathways such as an increase in KRAS signaling and epithelial to mesenchymal transition, as well as the NF-KB inflammatory pathway, were all upregulated in our more migratory populations **(Figure 8B).** These pathways correlate with an increase in cancer progression and would be expected to be found in more migratory cells. Interestingly, we also found that the ITGB4 pathway was also enriched in the migratory cells matching nicely with our protein expression flow cytometry data.

**Figure 8.**
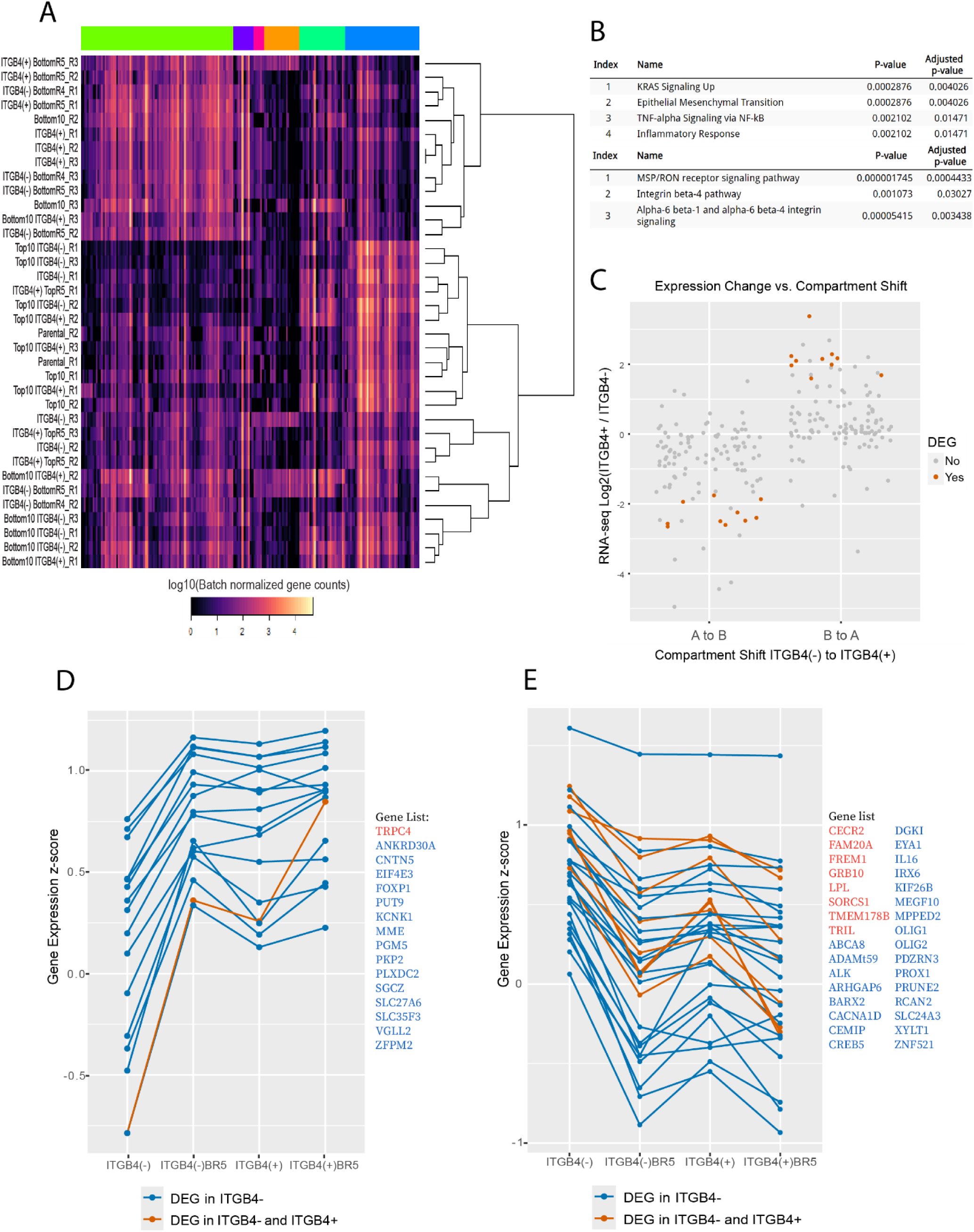
Gene expression changes partially correlate with compartment shifts following constricted migration. **A)** The top 500 most variable genes were used to cluster RNA-seq data across 10 conditions (3 replicates each). Light colored genes are more expressed while dark colored genes are less expressed. Top colorbar shows hierarchical clustering on the genes. The lime cluster represents genes upregulated in migratory cells while the blue cluster represents genes upregulated in the non-migratory cell conditions. **B)** Pathway enrichment analysis of the Lime cluster of genes (upregulated in migratory cells) using EnrichR (MSigDB Hallmark 2020 and NCI-Nature 2016). **C)** Gene expression changes from ITGB4(-) to (+) cells for the genes in ITGB4(-) to (+) compartment changes. Significantly differentially expressed genes are shown in orange, others grey. **D)** Gene expression Z-score across ITGB4 conditions for genes in the 250kb bins with positive compartment shift slopes (Fig 7A). Blue = differentially expressed in ITGB4(-) vs. (-)BR5. Orange = significantly differentially expressed in both ITGB4(+) and (-) cells after 5 rounds of migration. The gene list is provided to the right. **E)** Gene expression Z-score across ITGB4 conditions for genes in the 250kb bins with negative compartment shift slopes (Fig 7A). Colors as in (D).

Next, we asked whether these gene expression differences among cell populations corresponded to compartment shifts we had observed. We first took the list of genomic regions that switched compartments between ITGB4(-) and ITGB4(+) cell populations and looked at the RNA expression profiles of the genes within them in those same cell groups. While only a few of these genes were classified as significantly differently expressed, we observed that the level of RNA expression shifted in the direction expected based on the compartment shift (A to B showing downregulation in ITGB4(+) vs. ITGB4(-) and B to A showing upregulation; **Figure 8C**). We then examined genes in the regions that shifted following migration of ITGB4(-) and ITGB4(+) cells. We observed that for both B to A shifts **(Figure 8D)** and A to B shifts **(Figure 8E)**, genes that change expression move in the same direction as the compartment shifts (compare to **Figure 7A**). Only a subset (∼10%) of the genes found in these compartment shifting regions were differentially expressed, suggesting that compartment switching is not only for the purpose of facilitating gene regulation, but could also play a role in the physical state of the cell. However, those genes that did change were concordant with the direction of the compartment shift.

## Discussion

Many factors are involved in determining whether cancer cells will metastasize and what their phenotype will be once they reach a distant site. Here, we focused on one part of this complex process: the ability of A375 melanoma cells to migrate through constrictions smaller than their nucleus. Previous work had identified 3D genome structure changes and gene expression changes in melanoma cells that had migrated through 10 rounds of constriction and gained a highly invasive phenotype^10^, but it was unclear whether these differences arose from a pre-existing subpopulation selected by the rounds of migration or whether they were induced during the process of constriction. Here, our results have begun to disentangle this conundrum. We identified a cell surface protein, ITGB4, that was highly expressed across all our highly migratory sequentially constricted cells, and we found that ITGB4 was already expressed on a subpopulation of cells before migration. We found that sorting out high ITGB4-expressing cells from the parental A375 population yielded a subgroup of cells that were already as migratory as the cells that had passed through 10 rounds of constriction (Bottom10). This result, by itself, could mean that the “changes” we saw in sequentially constricted cells were simply the result of isolating this already highly migratory ITGB4 expressing population. However, further investigation revealed that there are key ways that sequentially constricted Bottom10 cells are different from this initial migratory ITGB4(+) subpopulation. Examining cellular elongation, behavior when embedded in 3D collagen matrices, and ability to migrate through even smaller pores, it became clear that this initial ITGB4(+) population is an intermediate between the non-migratory cells in the initial population and the highly invasive cells after sequential passage through constrictions. Therefore, detailed analysis and further migration of this ITGB4(+) subpopulation provided us a window into what genome architecture differences pre-exist and which are induced by sequential constriction.

ITGB4 expression by itself is not sufficient to explain why this initial subpopulation is more proficient at constricted migration, since knocking down ITGB4 resulted in only modest decreases in migration. Instead, this subpopulation contains 3D genome architecture and gene expression differences throughout the genome that could play a role in defining an initially increased migration ability. Indeed, both by genome compartment signatures and gene expression signatures, ITGB4(+) cells clustered with highly migratory cells, and genes defining these differential signatures were involved in metastasis and migration-relevant processes such as epithelial-mesenchymal transition. Whether we started from the entire parental A375 population, or a population grown from a single cell clone, we found that ITGB4(+) cells showed consistent regions of the genome with altered compartmentalization as compared to ITGB4(-) cells.

As with their cellular appearance and behavior, the genome structure state of ITGB4(+) cells is an intermediate between the parental population and the Bottom10 cells. Further changes to the 3D genome occur in these cells only after constricted migration. Our finding that many genome structure changes are shared whether initially ITGB4(+) or ITGB4(-) cells are subjected to several rounds of constricted migration suggests that there are indeed certain genomic regions that are particularly susceptible to change with passage through constrictions.

Interestingly, these further changes that happen to cells after constricted migration not only make the cells better at migrating through even smaller constrictions, but also remove any dependence on ITGB4 for migration. We propose that, in this way, ITGB4 expression is conceptually similar to training wheels on a bicycle. Initially, for an inexperienced rider, training wheels provide important assistance, and removing them would increase the likelihood that a rider would fall, just as removing ITGB4 from a cell that is not yet “experienced” with constricted migration results in somewhat decreased migratory ability. But, once a rider has undergone further training and experience, removing the training wheels would make little difference, just as removing ITGB4 from Bottom10 cells no longer influences migration rates.

Our results also reveal that the final state of cells after sequential constriction is not simply a matter of selecting cells that already express ITGB4 and then further remodeling them through the process of constriction. Within our populations grown from individual cell clones, which have reduced heterogeneity, we have examples where almost no cells expressed ITGB4 initially, but still, after 10 rounds of constriction, ITGB4 became highly expressed. By carefully tracking ITGB4 surface protein levels before and after each round of migration, we found that signatures of selection were indeed present: more ITGB4-expressing cells were found immediately after passage through pores than before. However, substantial increases in migration and ITGB4 expression required multiple rounds of constriction. Further, there were also cases where ITGB4 expression was increased during cell growth between rounds of migration, suggesting that passage through the constriction activates processes which can continue to change the cell phenotype even after that acute stress is over. This experiment also clarified that it is truly the constriction, not other factors in the migration process, that results in ITGB4 increase: cells passing through wide 12µm pores were exposed to the same fibronectin and growth media conditions, but did not show increases in ITGB4 expression over time.

Overall, our study reveals that 3D genome structure differences may be important both for encoding an initial highly migratory cell identity as well as for stabilizing the further alterations in cell phenotype that occur after constricted migration. This conclusion is in line with other work in the field that emphasizes the importance of genome spatial compartmentalization in defining cell fate and cell type.^19^ Further work will be needed to identify what triggers and pathways are activating increases in ITGB4 and causing other changes to the 3D genome structure. With the sets of regions identified here, future work can investigate what properties make a genomic region a hotspot susceptible to compartment change after constricted migration. One notable feature among the regions we identified is that gene poor, peripherally located chromosomes experienced more changes than gene rich internal chromosomes. Coinciding with this, it is important to note that while gene expression levels correlated with the compartment alterations we observed, only a small subset of genes experienced changes in expression. This suggests, as described previously^36,37^, that the migration-favoring 3D genome alterations can also be important to nucleus physical properties rather than only gene regulation.

We expect that the changes we observe here relate to the physical microenvironment-induced alterations in chromatin modifying enzymes that have been shown in other contexts.^11,12,22^ Indeed, initial analysis suggests that one common feature between the genomic regions that change compartment with constriction is their tendency to be marked with H3K27me3 (according to annotations in EnrichR). This suggests that the changes in this mark that can be induced by passage through constrictions^22^ could result in the long-term compartment alterations we observe here.

From a therapeutic perspective, our results indicate that there are likely to be RNA and 3D genome signatures within subpopulations of cells in a tumor that indicate a cell’s initial propensity to undergo migration. Such invasive gene expression signatures have previously been discussed^38,39^, but our study clarifies pre-existing 3D genome signatures and how these may further change during the process of cell migration. Our results also show that a whole suite of cellular and genomic changes accompany ITGB4 status, and knocking down ITGB4 only modestly reduces migration. Thus, our results also caution against using a single factor like ITGB4 as a prognostic marker or therapeutic target.

## Limitations of the study

When analyzing specific genomic regions that alter their gene expression or structure during a process like constricted cell migration, we are always faced with a tradeoff between the depth of information we can obtain, the precision of genomic localization, and the ability to track a cell in real time. In this study, we have used sub-population sorting to examine differences within a heterogeneous cell population. However, even within a population grown from a single cell clone or sorted based on certain characteristics, we know that there is still inherent heterogeneity and cell to cell variability that is not captured in our bulk Hi-C and RNA-seq analyses. Further, even when we do track individual cell properties such as ITGB4 surface protein expression, aspect ratio, or behavior in 3D collagen matrices, we are limited to cells at certain timepoints (before migration, after each round of migration) rather than tracking changes in an individual cell over time. Finally, we are examining a single melanoma cell line. Our past work suggests that some of the 3D genome structure alterations we observe are shared even among different cancer types^10^, but the specific signatures of migration ability may differ for cells from different patients or different tumor starting points.

## Acknowledgments

We thank all members of the McCord lab for helpful discussion and feedback. We thank Dibyayan Maity and Tehya Daniels for their help with early troubleshooting experiments. We acknowledge the UTK Advanced Microscopy and Imaging Center and Genomics Core for assistance with imaging and sequencing and Trevor Hancock for assistance with flow cytometry. We thank Chandler Ross for assistance with image analysis. This work was supported by the National Institutes of Health [NIGMS grant R35GM133557 to R.P.M].

## Author contributions

C.P. and R.P.M. designed the study. C.P., R.G., T.H.O., S.J.B., and A.S. performed experiments. J.H.G. and A.R.G. performed bioinformatic analyses. C.P. and R.P.M. wrote the paper with contributions from all authors.

## Data Availability

Raw and processed Hi-C and RNA-seq data files are available at GEO accession number GSE275397.

## Methods

### A375 Cell Culture and Transwell Migration Assays

A375 cells were obtained from ATCC (CRL-1619). Cells were verified to be negative for mycoplasma and were grown using complete DMEM medium (Corning-10-013-CV; 10% FBS, 1% Pen-Strep, 1% L-Glutamine) at 37°C supplied with 5% CO_2_.

Transwell filters with 5μm pore sizes (VWR-10769-236) and 3um pore sizes (Corning-353096) were used. Briefly, the bottom of each filter was coated with 40μl of 10 μg/ml fibronectin for ∼30 min. 24-well plates were prepared for Transwell migration assay by adding 500 μl of 1× DMEM (Corning) with full supplements per well. A375 cells were detached from culture dishes at 80–90% confluency and aliquoted to 100,000 cells per 100μl of unsupplemented 1×DMEM. Each Transwell was placed into its corresponding well of the 24-well plates, and 100μl of the cell suspension was added to the top of each filter. Cells were incubated at 37°C, 5% CO2 and allowed to migrate for 24hr.

After the 24hr incubation, migration efficiency was quantified as follows; first, freely floating cells were removed from the top of the filter (unmigrated cells) and from the well beneath the filter (migrated cells) and placed in two separate tubes. Then, 400μl of trypsin was added into the bottom chamber of the 24-well plates, and 200μl of trypsin was added into the top chamber to detach any remaining attached cells. Recovered cells after trypsinization were added to the unmigrated or migrated tubes, accordingly. Cells were spun down (1,000 rpm, 5 min) and resuspended in 100uL of DMEM. 15ul from each tube was collected and counted (using trypan blue) on a hemacytometer to calculate % migration such as: #bottom/(#top+#bottom).

For sequential rounds of migration, the migrated cells were seeded into a new well of a 24-well plate to expand. When cells reached 80–90% confluency, another Transwell migration was performed as previously described. Only the bottom cells were collected and continued to be used for sequential rounds of migration.

### Flow cytometry and Fluorescence Activated Cell Sorting ITGB4 Staining

All cytometry and FACS experiments followed the same protocol for ITGB4 staining. Cells were harvested using trypsin (T25, 6-well, 24-well) and spun at 1000rpm for 5 minutes. Supernatant was removed and cells were resuspended in 400uL of PBS containing 10% FBS and 1.25ug of Abcam FITC Anti-ITGB4 (ab22486). Cells were then incubated on ice for 30 minutes with slight agitation every 10 minutes (flicking) to ensure antibody distribution. Cells were then diluted with 350uL of PBS containing 10% FBS and mixed by pipetting before being spun at 1000rpm for 5 minutes. Supernatant was removed and cells were washed 2 more times with 400uL of PBS containing 10% FBS to ensure excess antibody was washed away. Unfixed cells were used for FACS, but for flow cytometry quantification, cells were then fixed by adding 500uL of 1% formaldehyde and incubated at RT for 10 minutes. Cells then were washed 2 times with 400uL of PBS containing 10% FBS. Finally, cells were suspended in 500uL of PBS containing 10% FBS and run on a flow cytometer.

### Flow Cytometry and FACS Analysis

For FACS experiments, cells were sorted by either a FACS Aria cell sorter (Hi-C replicate 1 for ITGB4(+) and ITGB4(-) cells, and some initial 5um migration experiments) or a Sony MA900 cell sorter (All other experiments). To determine the cutoff for ITGB4 expression, an Unstained cell population was used as a negative control. These cells followed the same staining protocol described above but did not include 1.25ug of the ITGB4 antibody. For the FACS Aria, once the cells were gated for the live population and doublets were removed, anything that expressed ITGB4 (FIT-C) above the Unstained condition was sorted into a (+) tube, while everything else was sorted into a (-) tube. To get a more stringent cutoff for ITGB4(+) and ITGB4(-), we re- sorted using the Sony 6000 sorter and after gating for live cells and removing doublets we again used an Unstained population of cells to get a cutoff, but then set 2 thresholds, one above and one below the Unstained cutoff to create 3 populations. The highly ITGB4 expressing cells were sorted into the (+) tubes, the very little to no ITGB4 expressing cells were sorted into the (-) tube, and everything in between was not collected (Figure 1D).

For flow cytometry experiments, cells were run on a BD LSR II (Clone ITGB4 expression levels, confluency and media depletion studies, and ITGB4(+) cells expression over time) or a Cytek Northern Lights flow cytometer (Sequential migration with flow and RNAi experiments). An Unstained population of cells was also used as a negative control to set a threshold value for ITGB4 expression. Cells were gated on real events, and doublets removed. To calculate the percentage of cells that express ITGB4, the percentage of events (cells) that had FITC (ITGB4) values over the Unstained condition were taken. Average ITGB4 intensity values are the average of the raw values of FITC collected by either flow cytometer for all cells in the condition. All analysis was done with Floreada.io (https://floreada.io).

### Scratch assay

ITGB4(+) and ITGB4(-) cells were grown in single wells of a tissue culture treated 6-well plate to high confluency of ∼90% in supplemented DMEM. Media was removed and each well was washed with 1mL of PBS. Scratch was made using a 10uL pipette tip by lightly applying pressure while moving along the bottom of the plate from bottom to top. Cells were then washed 3 times with unsupplemented DMEM to remove any debris. Finally, 1mL of DMEM supplemented with 0.1% FBS was added to the wells. Plates were then imaged for 24hr at 10min intervals using a 10x objective inside the live cell chamber of an EVOS II. Cells were kept at 37C, 5% CO2, and high humidity for the entirety of the image collection. Images were then stacked and turned into movies using Fiji Image Software.

### 3D Collagen Matrix

Experiments investigating the morphology and migration of A375 subpopulation of cells (ITGB4(+), ITGB4(-), and Bottom10) in 3D environment were performed using single cell suspension in 3D collagen matrices as previously described.^40^ Briefly, 75,000 cells/condition were added into collagen matrices (Rat Tail, Corning 354236) with a density of 2.32 mg/ml. The pH of the collagen gels was normalized to a neutral pH and gels were incubated at 37°C for 30 min to solidify. After the incubation, media with full supplements was supplied to the collagen gels. Cells were allowed to migrate through the collagen for 96hr before imaging.

### Confluence and Media Depletion Tests

To test the sensitivities of ITGB4 surface protein expression, Parental cells that had been through multiple rounds of migration were used. For Confluency, Parental cells that had made it through 4 rounds of migration were used. Cells from an 80% confluent 6-well plate were seeded at 15% and 50% density in 3mL of supplemented DMEM and allowed to proliferate for 24hr. The following day cells were ∼30% and 95% confluent, respectively, and were stained for ITGB4 surface expression and ran on a flow cytometer as previously described. For media influence, Parental cells through 4 rounds of migration were also used. Cells from an 80% confluent 6-well plate were seeded at 25% density in 3mL of supplemented DMEM in 2 wells and allowed to proliferate for 24hr. The following day, both wells had their media removed and replaced with either fresh supplemented DMEM, or media from a fully confluent well of a 6-well plate (depleted DMEM). The following day both wells were lifted and stained for ITGB4 surface protein expression and ran on a flow cytometer as previously described. For combination treatments, Parental cells through 8 rounds of migration were used. Combination treatments followed the same procedures as described above.

### ITGB4 RNAi

All supplies for siRNA experiments were ordered from Dharmacon. RNAi experiments were done following the manufacturers protocol in 24-well plates with final volumes of 500uL per well. Briefly, Human ITGB4 siRNA SMARTpool (L-008011-00-0005) and Non-targeting Pool (D- 001810-10-05) were resuspended in 1x siRNA Buffer (B-002000-UB-100) to a final concentration of 5uM. 4 Eppendorf tubes were needed to create both the siRNA and the non-targeting control vectors. In the RNAi tube, 5uL of 5uM ITGB4 siRNA was combined with 45uL of unsupplemented DMEM and mixed by pipetting and incubated at RT for 5 mins.

Concurrently, the Non-targeting tube had 5uL of non-targeting RNAi mixed with 45uL of unsupplemented DMEM and incubated at RT for 5 mins. Also, 2 Eppendorf tubes containing 1uL of DharmaFECT 1 Transfection Reagent (T-2001-01) and 49uL of unsupplemented DMEM were mixed and incubated at RT for 5 mins. Following the incubation, RNAi and Non-targeting tubes were combined with a DharmaFECT tube (100uL total) and mixed by pipetting before incubating at RT for 20 minutes. Then, 400uL of antibiotic free DMEM (supplemented with 10% FBS, and 1% L-Glutamine) was added to each tube (500uL total). RNAi and Non-targeting transfection medias were then placed into respective wells containing 20,000 ITGB4(+) or Bottom10 cells and allowed to proliferate for 48hrs. After 48hr this process was repeated and fresh RNAi and Non-targeting transfection medias were given to the cells and another 48hr of proliferation occurred (96hr total) before being collected and ran on a flow cytometer. Non-transfected controls for both ITGB4(+) and Bottom10 cells were used to verify no expression knockdown occurred in the Non-targeting experiments.

### Hi-C experiments

The first biological replicate of ITGB4 sorted cells was processed with the protocol described in (Golloshi et al., 2018). Briefly, 10 million cells were crosslinked with 1% formaldehyde for 10 min. Crosslinked cells were then suspended in lysis buffer to permeabilize the cell membrane and were dounce homogenized. Chromatin was then digested in-nucleus overnight using the DpnII restriction enzyme. Digested ends were filled in with biotin-dATP, and the blunt ends of interacting fragments were ligated together. DNA was then purified by phenol-chloroform extraction. For library preparation, the NEBNext Ultra II DNA Library prep kit (NEB) was used for libraries with size ranges from 200-400bp. End Prep, Adaptor Ligation, and PCR amplification reactions were carried out on bead-bound DNA libraries. All other Hi-C experiments were performed using the Arima-HiC+ Kit from Arima Genomics following the protocol for Mammalian Cell Lines (A160134 v01) and preparing the libraries using Arima recommendations for the NEBNext Ultra II DNA library prep kit (protocol version A16041v01).

Sequencing was performed either through GeneWiz or at the UTK Genomics core using either an Illumina HiSeq or a Novaseq 6000 with 150bp paired end reads.

### Hi-C data analysis

Sequencing reads were mapped to the human genome (hg19), filtered, and iteratively corrected by HiCPro (https://github.com/nservant/HiC-Pro). All Hi-C contact matrices were scaled to a sum of 1 million interactions to allow comparisons between conditions in downstream analysis. For library quality and mapping statistics, see Sup. Table 1.

Compartment analysis was performed by running principal component analysis using matrix2compartment.pl script in the cworld-dekker pipeline available on GitHub (https://github.com/dekkerlab/cworld-dekker). The PC1 value was then used to determine compartment identity for 250kb binned matrices. We considered PC1 values greater than 0.01 to be A compartment and less than −0.01 to be B compartment. To define compartment switches between conditions, PC1 values much go from positive to negative (A to B) or negative to positive (B to A). For compartment shifts between conditions, the difference between conditions had to have a PC1 value change greater than 0.04 (Towards A) or less than -0.04 (Towards B). This amount of PC1 change correlates with a 20% change in compartment score. For clustering analysis, the top 200 most variable bins were used for compartment profile clustering.

Slope determination for shifted bins was completed as follows: Conditions were ordered from least to most migratory (i.e., Top-Unmigrated-Bottom10) and their PC1 values for all 250kb bins were determined. For each bin, the slope of the data points from least to most migratory was calculated and the bins were sorted from highest to lowest slope. Standard deviation of all slopes was calculated, and bins that had slopes ±1.5 standard deviations from the mean were considered for further analysis. Positive slopes were considered bins that shifted more towards the A compartment, whereas negative slopes were considered bins that shifted more towards the B compartment.

To determine genes within shifted or switched bins, the bins of interest were overlapped with the UCSC Genes “knownGene” track within genome hg19 and extracted for further analysis.

### RNA-seq experiments

For all RNA-sequencing samples, RNA was extracted using the Qiagen RNAeasy plus mini kit. In short, cells were lysed and homogenized, spun down in gDNA eliminator columns to remove any genomic DNA. All samples were washed with ethanol, and the total RNA was eluted. The quality and quantity of the RNA were quantified using a nanodrop. Total RNA was then sent for library preparation (polyA mRNA approach) and sequencing at GeneWiz.

### RNA-seq analysis

The fastq reads were first processed with BBDuk tool (https://github.com/kbaseapps/BBTools), performing adapter trimming. Adapter trimmed reads were processed for quality trimming using the BBDuk tool to discard reads with quality score lower than 28. Following the adapter and quality trimming steps, the reads were aligned to the reference genome hg19 using STAR aligner (https://github.com/alexdobin/STAR). Finally, the mapped reads were sorted based on genomic coordinates and feature count was performed with HTSeq-Counts (https://github.com/simon-anders/htseq). Differential gene expression was determined using DESeq through Galaxy. RNA-seq profile clustering was done by taking the 500 most variable genes throughout all conditions.

## Supplementary Figures

**Sup. Figure S1.**
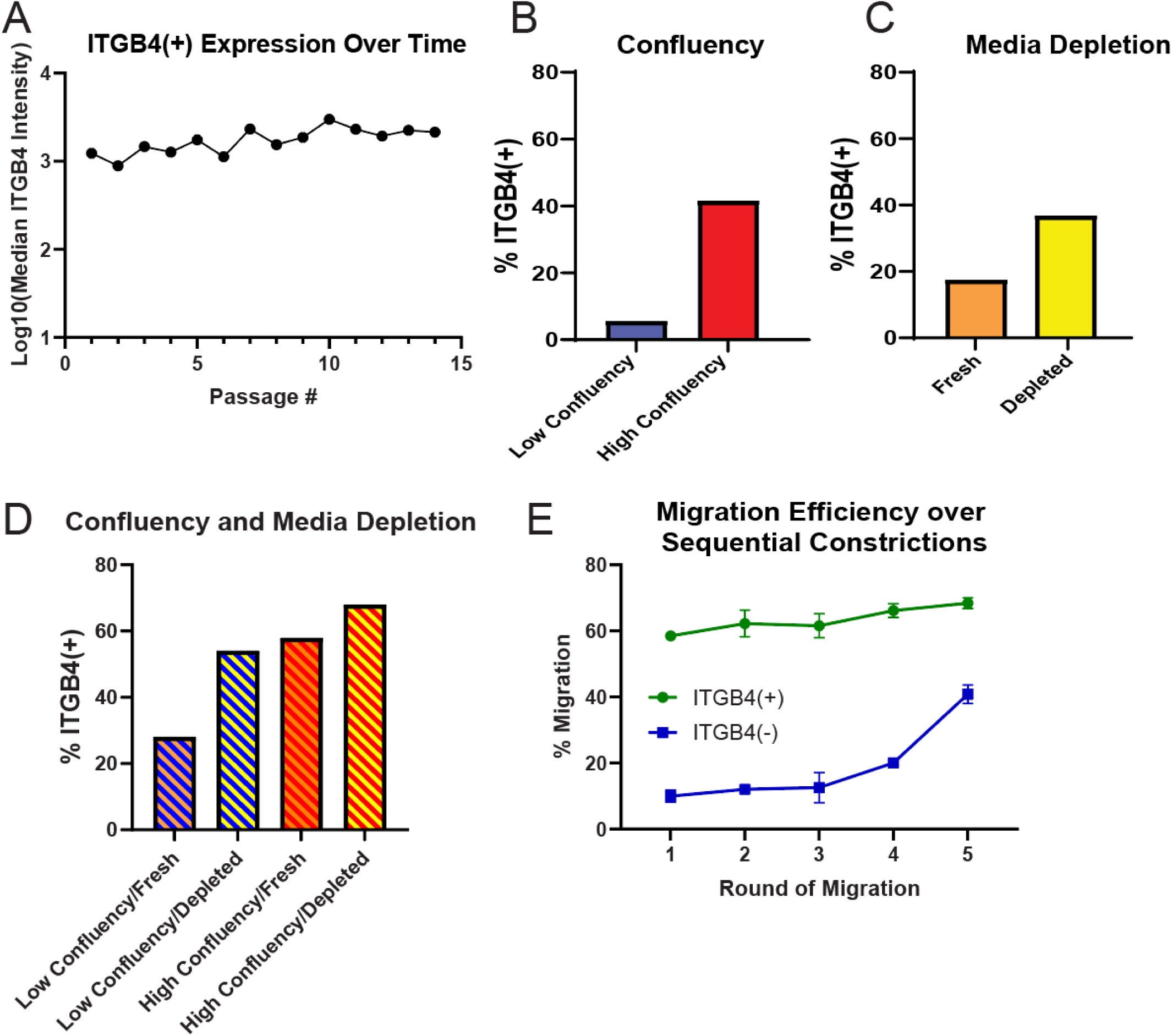
ITGB4 is sensitive to outside parameters. **A)** The ITGB4 intensity is measured by fluorescent antibody label in flow cytometry for the sorted ITGB4(+) population as cells are cultured across subsequent passages. The Log10 of the median of the ITGB4 intensity distribution is plotted for each timepoint and remains high across subsequent passages. **B)** The percentage of A375 cells expressing ITGB4 (measured by flow cytometry, gated based on negative control) when cells are at low confluency (30%) and high confluency (90%). Cells were taken after 4 rounds of migration and seeded for high or low confluency. **C)** The percentage of A375 cells expressing ITGB4 when cells are given freshly made media with full nutrient levels compared to when cells were given depleted media from 100% confluent A375 cells. Cells taken after 4 rounds of migration and seeded in different media conditions. **D)** The percent of A375 cells expressing ITGB4 when exposed to a combination of conditions from B and C. Cells were taken after 8 rounds of migration for exposure to different conditions. **E)** Sorted ITGB4(+) and (-) cell populations were subjected to 5 rounds of migration through 5µm Transwell pores. The percentage of cells that passed through the pores in each round is shown.

**Sup. Figure S2.**
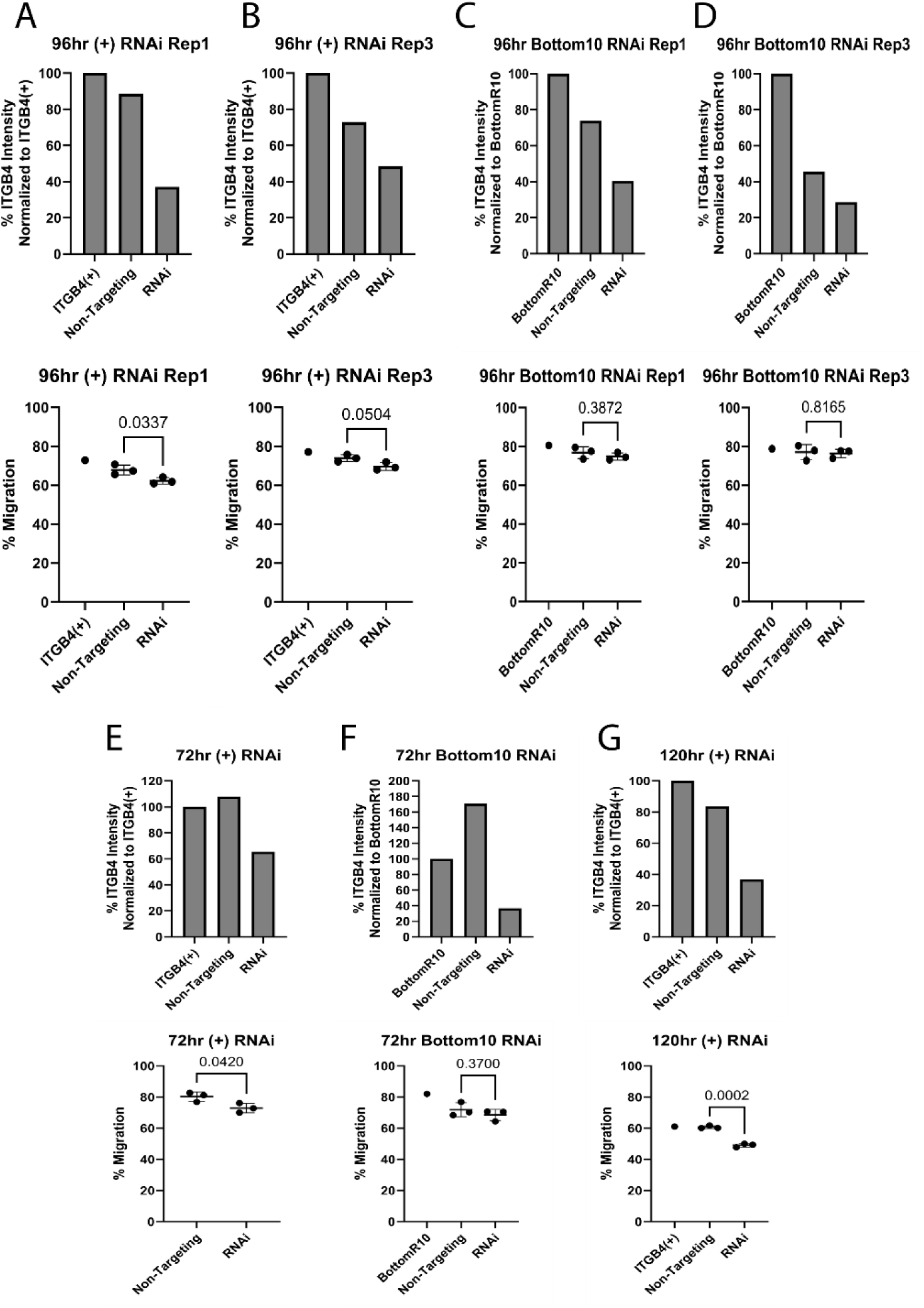
Extra ITGB4 RNAi replicates show similar results for 96hrs of treatment. **A, B** Replicates 1 and 3 of ITGB4(+) cells given ITGB4 RNAi. Top graph shows the amount of ITGB4 expressions for untreated, Non-targeting Control, and RNAi treatments. The bottom graph shows the respective migration percentage through 5µm Transwell pores. All replicates showed a reduction in migration ability. **C, D** Replicates 1 and 3 of Bottom10 cells given ITGB4 RNAi. Top graph shows the amount of ITGB4 expressions for untreated, Non-targeting Control, and RNAi treatments. The bottom graph shows the respective migration percentage through 5µm Transwell pores. All replicates showed no change in migration ability following loss of ITGB4 expression. **E** A single replicate of ITGB4(+) cells given RNAi for 72hrs straight. The top graph shows ITGB4 expression where the bottom graph shows migration ability for each condition. (ITGB4(+) control not collected due to technical error). 72hr did not give as reproducible a knockdown as 96hr. **F** A single replicate of Bottom10 cells given RNAi for 72hrs straight. The top graph shows ITGB4 expression where the bottom graph shows migration ability for each condition. At 72hr we saw no difference in migration ability. **G** A single replicate of ITGB4(+) cells given RNAi for 120hrs (48+72). The top graph shows ITGB4 expression where the bottom graph shows migration ability for each condition. We saw a strong knockdown and migration difference, but the cells were less viable after 120hrs of treatment.

**Sup. Figure S3.**
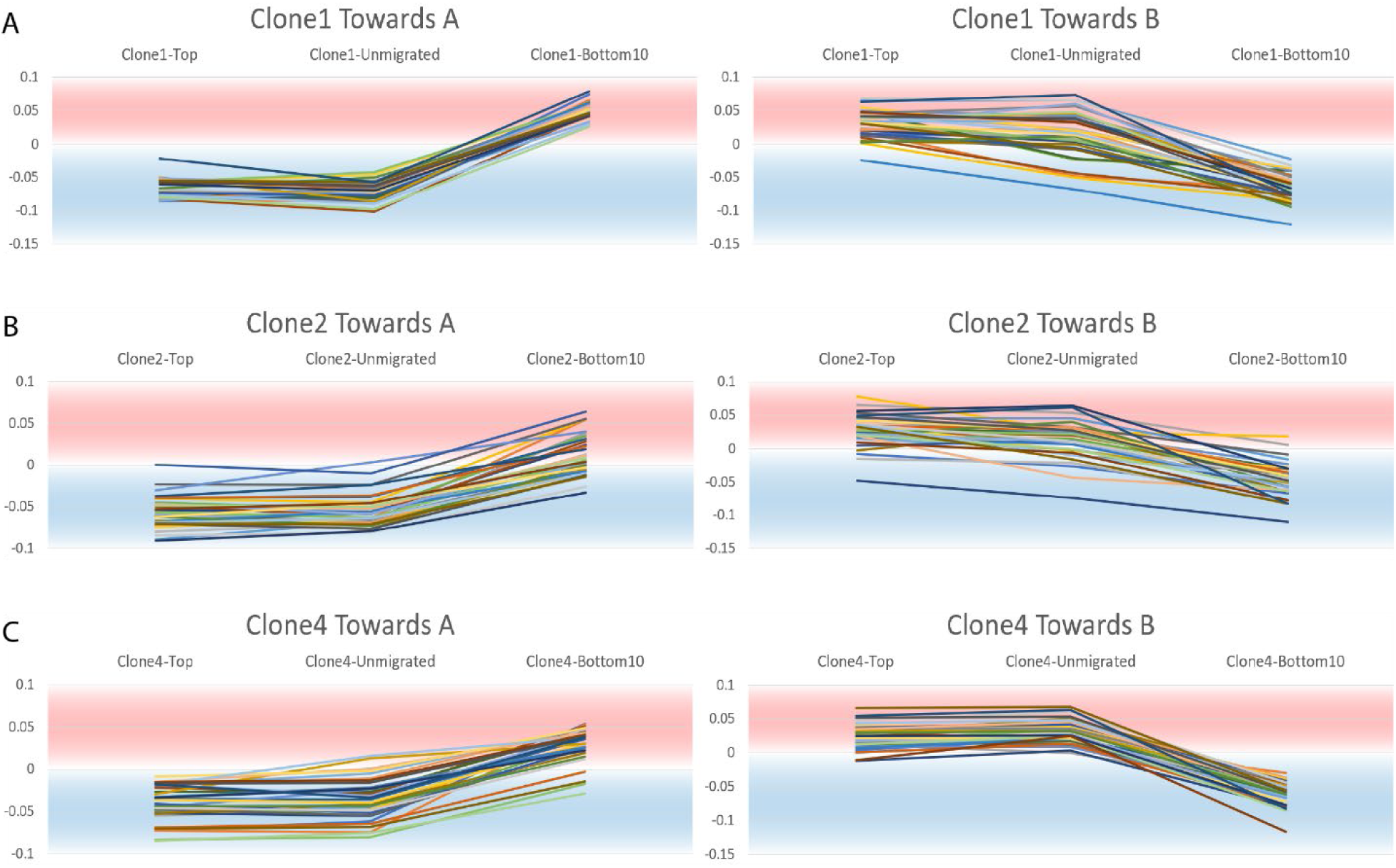
All Clonal populations had genomic regions that shift after constricted migrations. **A-C** The top 30 250kb binned regions with positive slopes (Towards A) and negative slopes (Towards B) as cells increase migration ability for Clone1, Clone2, and Clone4 cells. Each bin’s compartment value is plotted for Top10, Unmigrated, and Bottom10 cells and tracked from left to right.

**Sup. Table S1.**
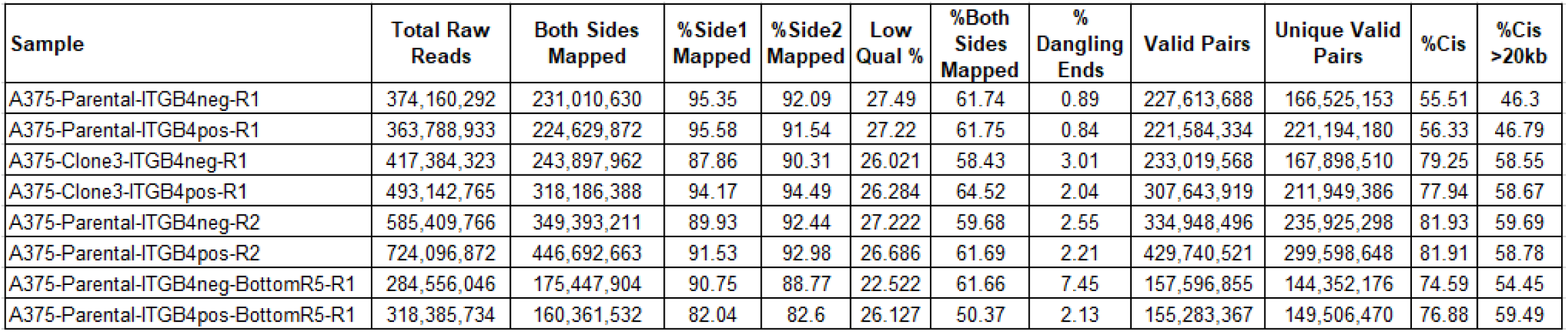
Sequencing and mapping statistics for all Hi-C datasets in this study.

## Movies

Movie 1: 24hr scratch assay of ITGB4(-) cells. Images taken every 10 minutes at 10x. Scale bar = 300µm.

Movie 2: 24hr scratch assay of ITGB4(+) cells. Images taken every 10 minutes at 10x. Scale bar = 300µm.

Movie 3: ITGB4(-) cells embedded in 3D collagen matrix. Cells acclimated for 96hr before 24hr movie was taken. Images taken every 8 minutes at 20x. Scale bar = 100µm.

Movie 4: ITGB4(-) cells embedded in 3D collagen matrix. Cells acclimated for 96hr before 24hr movie was taken. Images taken every 8 minutes at 20x. Scale bar = 100µm.

Movie 5: ITGB4(+) cells embedded in 3D collagen matrix. Cells acclimated for 96hr before 24hr movie was taken. Images taken every 8 minutes at 20x. Scale bar = 100µm.

Movie 6: ITGB4(+) cells embedded in 3D collagen matrix. Cells acclimated for 96hr before 24hr movie was taken. Images taken every 8 minutes at 20x. Scale bar = 100µm.

Movie 7: A375 Bottom10 cells embedded in 3D collagen matrix. Cells acclimated for 96hr before 24hr movie was taken. Images taken every 8 minutes at 40x. Scale bar = 70µm.

